# Hypoxia inhibits newt skeletal muscle dedifferentiation

**DOI:** 10.1101/2024.11.29.626015

**Authors:** Georgii Vdovin, Hannah E. Walters, Konstantin E. Troyanovskiy, Yugo Tsuchiya, Maximina H Yun

## Abstract

Unlike mammals, newts are able to regenerate lost muscle tissue by dedifferentiating postmitotic muscle cells, which re-enter the cell cycle to form myogenic progenitors. This ability is considered central to the process of limb regeneration, yet the mechanisms underlying it remain poorly understood. Here, we leverage high-resolution time course bulk transcriptomics profiling to characterise molecular changes during differentiation and dedifferentiation of newt myotubes. We uncover that myogenesis is accompanied by a metabolic shift from glycolysis to OXPHOS, which is partially reverted upon dedifferentiation. By analysing early events during dedifferentiation, we identify TGFβ and members of the oxygen-responsive HIF family as putative plasticity inducers. We detect a stark, transient upregulation of HIF3A early during dedifferentiation, followed by a gradual downregulation of HIF2A. Finally, we show that hypoxia inhibits myotube cell cycle re-entry. These data provide a valuable resource to identify regulators of cellular plasticity and offer insights into the impact of oxygen signalling in regenerative processes.

## INTRODUCTION

Upon limb amputation, salamanders such as axolotls and newts are capable of proliferation-driven epimorphic regeneration, which involves the formation of a mass of undifferentiated cells with restricted potential known as a blastema (Kragl et al., 2009). This regenerative ability remains intact from the larval stage and extends well into adulthood (Gómez and Echeverri, 2021). There are two main cellular sources for the formation of blastema which are thought to be used in different, yet not completely understood proportions among salamanders. The animals either draw from pre-existing stem cell pools such as Pax7^+^ satellite cells in muscle, or generate regenerative progenitors from differentiated cell types via dedifferentiation, with each cell type typically contributing to its respective progenitor cell pool (Brockes and Gates, 2014; Kumar et al., 2000; Sandoval-Guzmán et al., 2014; Wu et al., 2015). Recent findings suggest that newts undergo a developmental switch from one mode of regeneration to another as they undergo metamorphosis (Tanaka et al., 2016; Yu et al., 2022), with adult newts adopting dedifferentiation as a mechanism of muscle regeneration.

*Notophthalmus viridescens* (the Eastern newt) constitutes an excellent model to study this regenerative process, with an established newt limb-derived A1 cell line (Ferretti and Brockes, 1988) that can be induced to differentiate and dedifferentiate to and from myotubes (Tanaka et al JCB 1997). Upon serum deprivation, A1 cells differentiate into polynucleated myotubes which express key muscle markers. In response to re-exposure to normal serum levels, these myotubes re-enter the cell cycle and dedifferentiate, in stark contrast to terminally differentiated mammalian myotubes (Wang and Simon, 2016; Yun et al., 2014). At the cellular level, myotube dedifferentiation relies on sustained ERK activation (Yun et al., 2014), p53 downregulation (Yun et al., 2013) and pRB phosphorylation (Tanaka et al., 1997). The serum factors required for eliciting this striking response constitute, at least in part, bone morphogenetic proteins (BMPs) which are cleaved by serum proteases (Wagner et al., 2017). Myotube dedifferentiation can be further elicited by paracrine signals derived from other cell types, including senescent cells (Walters et al., 2023). Still, we currently lack a comprehensive picture of the stimuli required for dedifferentiation as well as the molecular network underlying this remarkable cellular phenomenon. Importantly, *in vitro* generated myotubes recapitulate endogenous muscle fibre behaviour when implanted in newt limb blastemas (Lo et al., 1993), and key insights obtained using the A1 model have been validated *in vivo* (Wagner et al., 2017; Walters et al., 2023), highlighting the relevance of this system for deepening our knowledge of the limited reprogramming transitions which are critical for complex tissue regeneration. Understanding how newts are able to modulate cellular plasticity stands to provide useful insights for regenerative medicine, both in the context of muscle injury, as well as age-related muscle loss known as sarcopenia (Domingues-Faria et al., 2016).

During muscle differentiation, proliferative stem cells and myoblasts adjust their energy metabolism from glycolysis to oxidative phosphorylation (OXPHOS) (Kierans and Taylor, 2021; Sin et al., 2015). This switch is regulated by oxygen availability and the HIF1A transcription factor, with hypoxia (oxygen <2% for mammalian muscle) preserving stemness of muscle stem cells (Endo et al., 2024) and supporting stem and highly proliferative cells overall, including cancer (Chen et al., 2023; Das et al., 2010; Forristal et al., 2010; Kim et al., 2006; Mohyeldin et al., 2010; Pircher et al., 2021; Sandvig et al., 2017).

More broadly, cellular response to hypoxia is mediated by three genes termed hypoxia-inducible factors: HIF1A, HIF2A and HIF3A (Webb et al., 2009). HIFα subunits are constitutively produced, but are hydroxylated by prolyl hydroxylase domain proteins (PHDs) under physiological oxygen levels (normoxia). Hydroxylation targets HIFs towards conjugation with E3 ubiquitin ligase member von Hippel-Lindau protein (VHL), poly-ubiquitylation and rapid proteasomal degradation. Under hypoxia, hydroxylation of HIFs is inhibited, upon which the alpha subunits stabilize, translocate into the nucleus and form a heterodimer with an oxygen-insensitive HIF1B/ARNT subunit. Together, they recruit the coactivators P300/CBP and attach to the hypoxia response element (HRE) sequence within the promoters of target genes, thereby regulating their expression (Kenneth and Rocha, 2008; Ravenna et al., 2016). All HIFs have structural similarity and all respond to oxygen levels, but perform distinct, context-dependent functions. While HIF1A is responsible for metabolic reprogramming, the functions of HIF2A / EPAS are less well-understood, though it has been linked to extracellular matrix (ECM) remodelling, erythropoiesis and angiogenesis (Gruber et al., 2007; Patel and Simon, 2008; Rankin et al., 2007). HIF3A is the most structurally distant and has no known role in muscle development, but is thought to inhibit both HIF1A and HIF2A by competitive binding to HIF1B subunit (Duan, 2016; Ravenna et al., 2016).

Interestingly, previous work suggests that oxygen signalling plays an important role in regenerative processes. The hypoxia sensor HIF1A has been shown to promote metabolic adjustments and stem cell enhancement (DeFrates et al., 2021; Zhang et al., 2015). Further, oxygen levels modulate the extent of bone formation and mineralisation in a dynamic manner, and hypoxic events are critical during the degradation and blastema formation phases during mouse digit tip regeneration (Sammarco et al., 2014).In a salamander regeneration context, chemical induction of hypoxia with CoCl₂ disrupts axolotl tail regrowth when applied within the first 30 minutes post-amputation, but has no significant effect if applied after wound closure (Baddar et al., 2021). Lastly, Jopling et al., 2012 showed enhanced dedifferentiation of the hypoxic zebrafish myocardium mediated by HIF1A upregulation. Yet, the impact of oxygen levels on skeletal muscle dedifferentiation remains unknown.

Here, we employ time-resolved transcriptomics on the newt A1 model of differentiation and dedifferentiation. We confirm the myoblast nature of the A1 cells and identify early molecular regulators of dedifferentiation including TGFβ and oxygen-responsive genes HIF3A and HIF2A. We uncover negative regulation of hypoxia signalling as a putative driver of muscle plasticity and show that a hypoxic environment curtails muscle dedifferentiation, underscoring the relevance of oxygen as a critical regulator of cell fate transitions.

## RESULTS

### Single-cell and bulk transcriptomics confirm the myoblast identity of newt A1 cells

The A1 newt myotube model has provided critical insights into mechanisms of dedifferentiation (Wagner et al, Dev Cell 2017, Yun et al PNAS 2013). The A1 cell line was derived from limb mesenchyme cells which exhibited indefinite proliferative capacity and myogenic potential (P. Ferretti and Brockes, 1988; Lo et al., 1993; Walters et al., 2023). However, its cellular identity had not been previously addressed.

To determine the identity and heterogeneity of the A1 cell line and thus confirm its physiological relevance to identify regulators of dedifferentiation, we performed single-cell RNAseq of mononucleated A1 cells. Single-cell data formed eight clusters, seven of which were closely related in space, while the remaining cluster (4) was more distinct at the transcriptomic level (Fig. 1A). The latter showed high expression of genes associated with protein synthesis and degradation (Fig. 1B), even when controlling for cell cycle effects (Supplementary Fig. 1A). The clusters were labelled as cells with putatively different protein synthesis rates, which were otherwise homogeneous. In both bulk and SC data, we identified high expression of smooth muscle markers such as alpha-smooth muscle actin 2 (Acta2) and transgelin (Tagln), as well as the mesenchymal marker vimentin (Fig. 1D). To further ascertain the identify of A1 cells, we compared expression profiles of A1 bulk RNAseq with those of mouse C2C12 myoblasts. This allowed us to capture expression of lowly expressed transcription factors generally not picked up by single-cell sequencing and achieve a more complete marker profiling (Fig. 1D&E). Notably, C2C12 myoblasts also exhibit a high level of smooth muscle marker gene expression (Fig. 1E) and vimentin, much like A1 cells. Importantly, A1 and C2C12 cells showed comparable expression of the key myoblast markers, including the myogenic factor 5 (Myf5) and Pax7. Together, these findings indicate that the A1 cell line is a largely homogeneous newt myoblast line.

**Figure 1.**
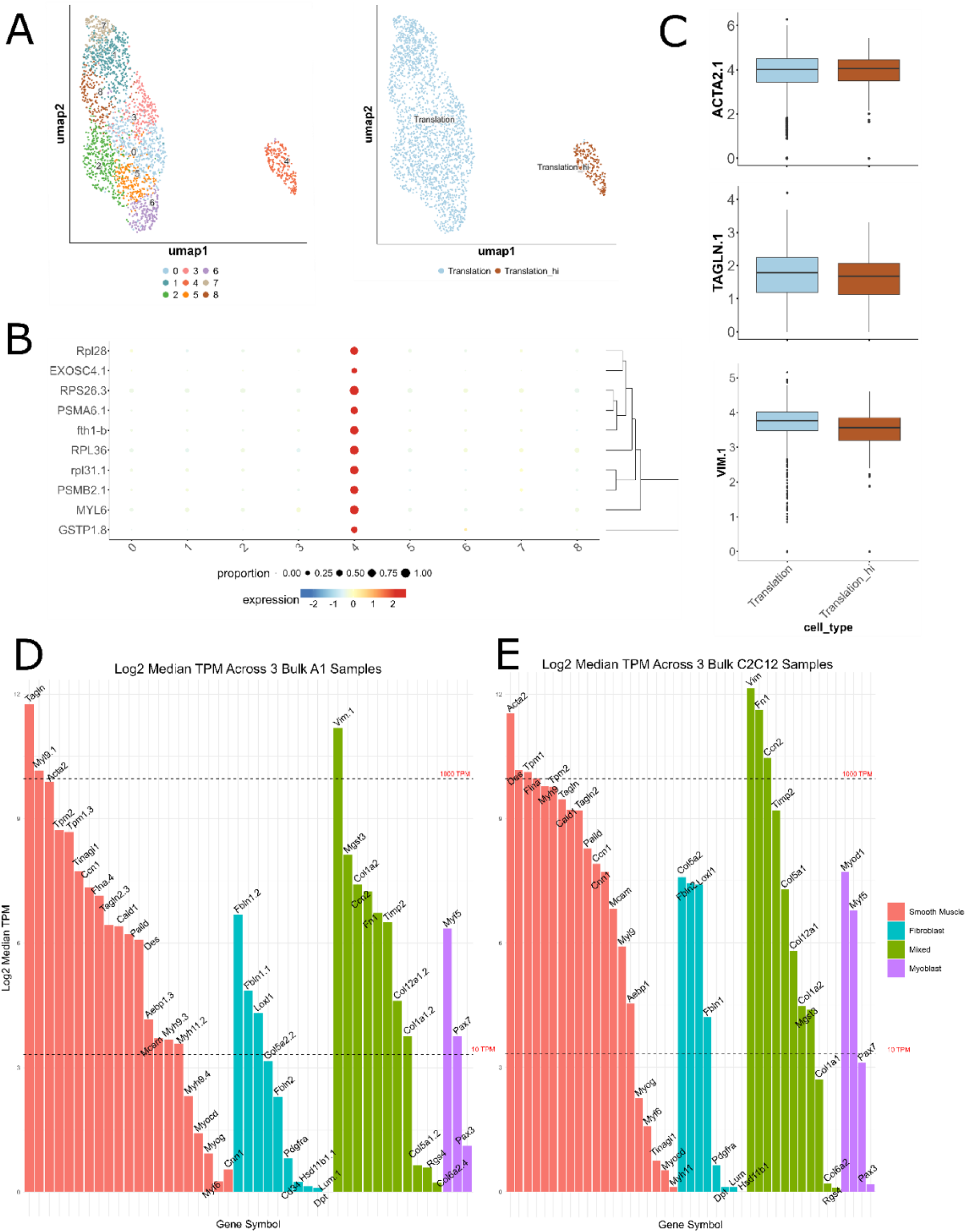
Heterogeneity and cell type identity of A1 cells. A) UMAP of single-cell transcriptomes of cultured A1 cells with Seurat-identified clusters (left), and biologically meaningful cell states (right). B) Bubble plot of top 10 annotated genes of cluster 4. C) Boxplot showing expression level of highest expressed genes associated with specific cell types, separately for two biological conditions. Top to bottom: smooth muscle actin, transglein and vimentin. D) Barplots depicting expression level of cell-type specific genes in A1 bulk RNAseq samples. Values are log2-transformed median transcript per million (TPM) across 3 replicates. Each unit change on the y-axis corresponds to a 2-fold change in expression level. Dotted lines indicate 10 TPM (bottom, borderline expression) and 1000 TPM (top, very high expression) marks. Colours indicate cell type with which a particular gene marker is associated. Myod1 was not found in the newt transcriptome annotation. E) Values for the same genes as in D, but for mouse C2C12 myoblasts. Note the similarity between A1 and C2C12 expression profiles.

### Time-resolved transcriptomics offers insights into myotube differentiation and dedifferentiation

We next set out to gain insights into the transcriptomic changes underlying differentiation from myoblasts into myotubes, and myotube dedifferentiation. Myotube formation was induced by seeding A1 myoblasts at confluence followed by serum withdrawal (to 0.25% FCS for 5 days). Myotube dedifferentiation was elicited by switching myotubes to a high serum medium (10%FCS), as previously described (Tanaka et al., 1997; Yun et al., 2014). To select optimal time points for transcriptomic profiling, we first precisely defined the timing of cell cycle exit and re-entry during differentiation and dedifferentiation. EdU-incorporation assays showed that serum deprivation, used to induce differentiation, reduced the proportion of cells in S-phase, with the number of cycling cells halving between 48 and 72 hours and reaching its lowest point at 120 hours (5 days; Fig. 2A). As expected, the change to a high-serum medium resulted in an increase in cycling mononucleates between 24 and 48-hours post-induction (Fig. 2A). In contrast, myotubes (MT) re-entered the cell cycle only 48 hours after dedifferentiation induction, reaching an average of ∼42% of cycling MT nuclei after 3 days in dedifferentiation medium (Fig. 2B). This data was used as a reference to choose the time points for bulk RNAseq, where early transcriptomic changes could be expected: 8h, 72h, and 120h (8h, 3 days and 5 days, respectively) for early and late differentiation, and 128h, 144h, and 168h (5 days + 8h, 6 days, and 7 days, respectively) for early and late dedifferentiation.

**Figure 2.**
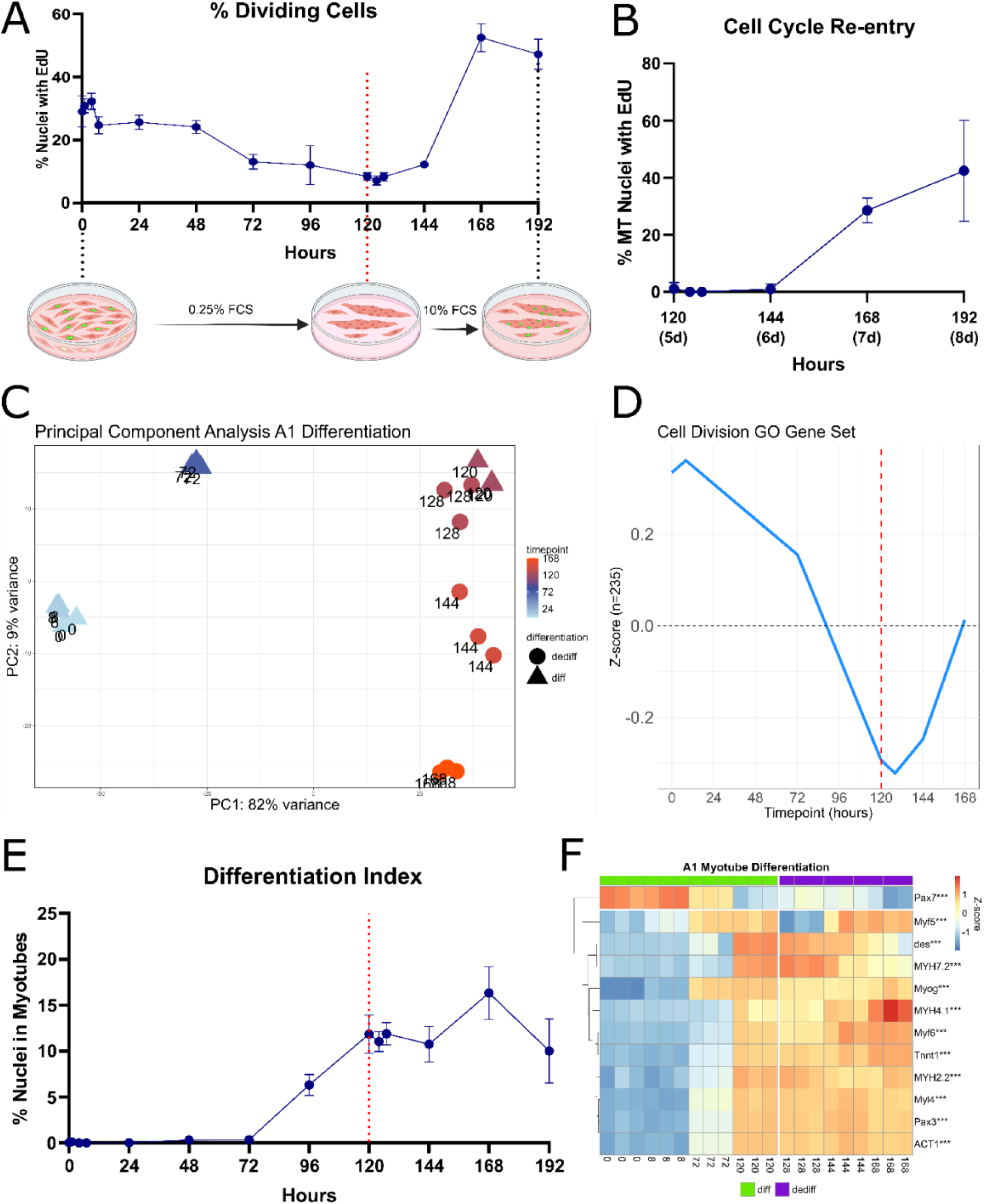
Cell cycle re-entry assay and myogenic differentiation time course of A1 cells. A) Percent of cycling cells across the time course as measured by percent of EdU-stained nuclei per time point, with experiment schematic shown below. Red dotted line in A, D, E - induction of dedifferentiation. Error bars - SEM of Y-values from four biological replicates. B) Cell cycle re-entry, calculated as a portion of EdU-incorporating myotube nuclei (Y-axis). Error bars - SEM of Y-values from four biological replicates. Hours after differentiation onset indicated on the X-axis. C) Principal component analysis on 500 most variable genes from the differentiation time course. Colour indicates time point, shape indicates differentiation conditions. Values derived from variance-stabilized count matrix. D) Gene set plot connecting the average Z-scores of all (235) genes annotated with Cell Division Gene Ontology term (GO:0051301) per time point. E) Differentiation index as measured by percent of nuclei in myotubes per time point. Error bars - SEM of Y-values from four biological replicates. F) Clustered heatmap of muscle differentiation markers. Time points are arranged in columns in chronological order, vertical cut and colour legend indicate induction of dedifferentiation. Stars on rownames indicate significance padj, after a Wald test comparing 120h to 0h (*p < 0.05, **p < 0.01 and ***p < 0.001).

Differentiating cells and dedifferentiating myotubes were harvested for transcriptomic profiling to capture time-dependent changes in gene expression. Of note, myotubes were purified ahead of profiling (from 120h onwards). Principal component analysis (PCA) showed separation by differentiation stage on PC1, while PC2 showed separation by serum exposure. Cells began reverting to their original transcriptional landscape and identity at 48 hours after dedifferentiation induction (168h; Fig. 2C). Notably, 48-hour dedifferentiating myotubes formed a distinct state from control cycling cells on PC2. The expression pattern of cell cycle-associated genes correlated with the observed dynamics of the cell cycle re-entry assay (Fig. 2A), reaching a low point in 5-day myotubes and rising again during dedifferentiation (Fig. 2D).

As expected, muscle differentiation and identity genes were among the top differentially expressed genes (Fig. 2F). The early myogenic marker Myf5 was retained during dedifferentiation, as previously described (Wang and Simon, 2016). However, we observed a surprising, transient downregulation of Myf5 at the 128h mark consistent across replicates, which returned to baseline levels within 24 hours. Importantly, a number of muscle marker genes (Desmin, MYH7, Myogenin) were downregulated upon dedifferentiation induction. Of note, some key myogenesis genes were initially absent from the transcriptome assembly but were added from aligning the data to an older assembly (Abdullayev et al., 2013; see Transcriptome Annotation under Methods). Myod1 was absent from all available *N. viridescens* assemblies. The acquisition and loss of muscle marker expression aligned with A1 cell differentiation dynamics (Fig. 2E), confirming the expected trends and overall dataset quality.

### Pathway analysis and metabolic changes of myogenic differentiation and dedifferentiation

Leveraging the temporal resolution offered by time course transcriptomics, we sought to explore the dynamics of pathway activity to identify potential drivers of differentiation and dedifferentiation. TGFβ signalling consistently emerged as the most significant and prominent upstream regulator (Supplementary Fig. 1B-D) and pathway (Fig. 3A) influencing these processes in both directions. Ingenuity pathway analysis (IPA) revealed that the TGFβ pathway was suppressed early during differentiation (as early as 8 hours after serum starvation), and rapidly reactivated 8 hours after induction of dedifferentiation. By 72 hours of serum starvation, its expression stabilized, aligning with the timing of cell cycle exit and the onset of myogenic differentiation (Fig. 2A&E). These processes appeared diametrically opposed, with TGFB1 playing a central role in both differentiation and dedifferentiation induction (Supplementary Fig. 1B-D).

**Figure 3.**
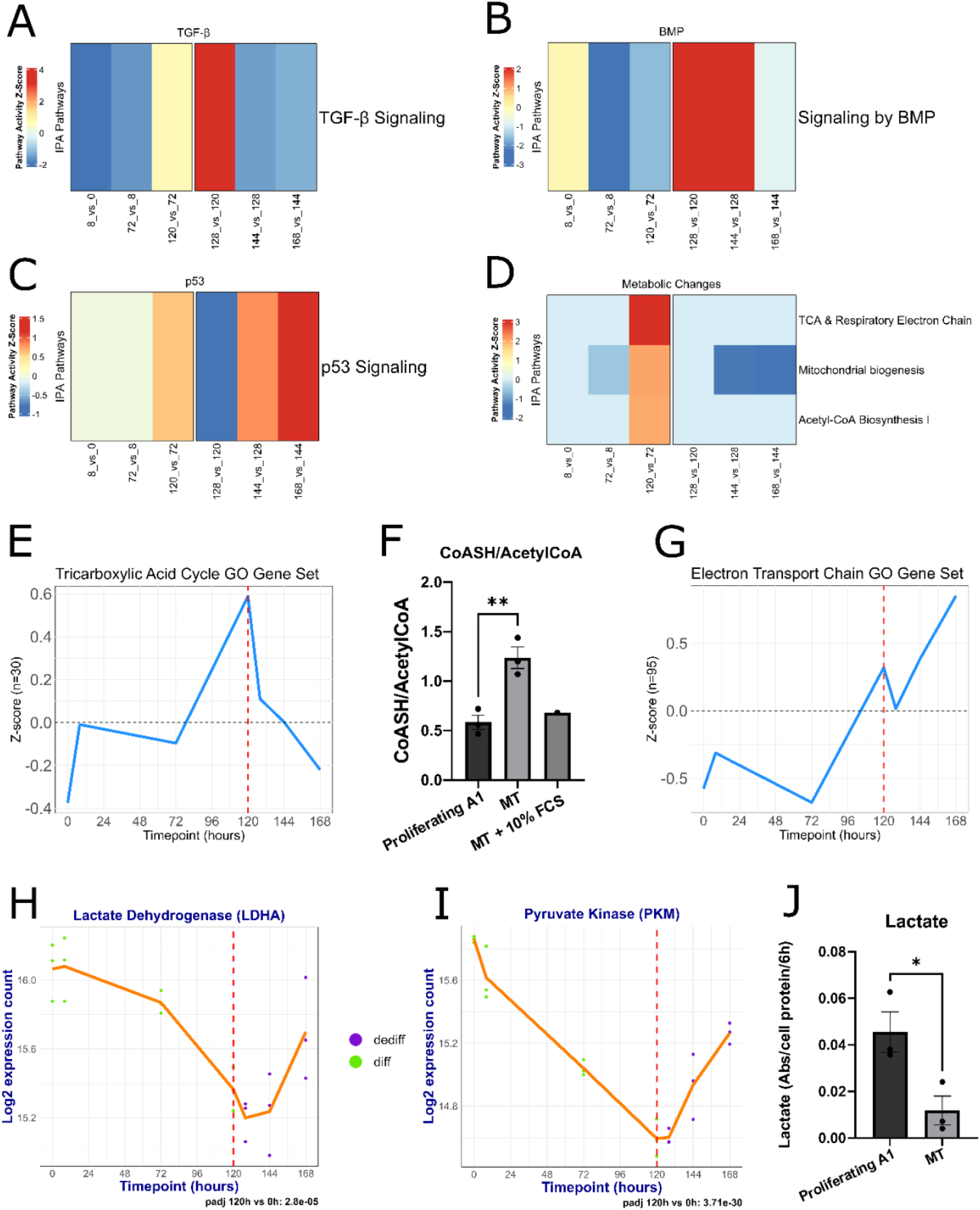
Pathway analysis and metabolic changes of myogenic differentiation and dedifferentiation. A-D) Heatmaps of pathway activity derived from Ingenuity Pathway Analysis. Shown in order: TGFβ, BMP, p53 and changes in energy consumption and respiration. Each cell represents a consecutive comparison to the previous timepoint, arranged in columns. Z-scores were taken directly from IPA. E) Gene set plot, connecting the average Z-scores of all (30) genes annotated with TCA (GO:0006099) gene ontology term. F) Ratio of free coenzyme A / Acetyl-CoA during differentiation and dedifferentiation of A1 myotubes. P-values here and in J are shown for an unpaired t-test (*p < 0.05 and **p < 0.01). G) Same as in E, but for 95 Electron Transport Chain gene set (GO:0022900). H, I) Gene expression changes of glycolytic enzymes Lactate Dehydrogenase A (H) and Pyruvate Kinase M (I) throughout the differentiation and dedifferentiation time course. Y-axis: log2-transformed normalized counts. X-axis - time point in hours. The orange line connects means of log2 counts per time point. Red dotted line and colour legend indicate induction of dedifferentiation. Adjusted pvalues shown after a Wald test comparing 120h to 0h. J) Changes in lactate production of undifferentiated vs differentiated A1 cells.

Inhibition of the TGFβ superfamily subgroup member BMP has been shown to reduce or block myotube cell cycle re-entry both in vitro and in vivo (Wagner et al., 2017), as well as to promote myogenesis in newt (Walters et al., 2023) and mouse C2C12 myoblast (Horbelt et al., 2015). We observed similar dynamics: BMP signalling downregulation took place from 72-hours during myotube differentiation (Fig. 3B), while its upregulation transiently occurred during dedifferentiation onset, starting as early as 8 hours post-initiation and persisting for 24-48 hours (6d-7d, Fig. 3B). Interestingly, the predicted changes in TGFβ activity preceded those of BMP in both differentiation and dedifferentiation, as supported by accompanying transient overexpression of pathway-associated SMADs and TGFβ targets RUNX2 and fibronectin (Lévesque et al., 2007) (Supplementary Fig. 3A-C), suggesting that TGFβ changes occur earlier than BMP. This overexpression is transient, and its detection was enabled by the high temporal resolution of our datasets.

Tight control of the p53 pathway is required for regeneration, particularly during muscle repair. Previous studies have demonstrated that p53 activation is necessary for differentiation and cell cycle exit, while premature stabilization of p53 disrupts differentiation. As regeneration progresses, p53 levels rise again, promoting the re-specialization of cells to restore the original limb structure (Yun et al., 2013). IPA precisely reflected the previously described changes, predicting p53 signalling activation at the 5d differentiated stage, with no changes in activity until then (Fig. 3C). Following dedifferentiation initiation, p53 was downregulated as early as 8 hours, before reactivation 24 hours later, consistent with published data.

During myogenic differentiation, myoblasts shift their energy metabolism from anaerobic glycolysis to oxidative phosphorylation in differentiated muscle (Elkalaf et al., 2013; Leary et al., 1998; Lyons et al., 2004; Shintaku et al., 2016; Sin et al., 2015), which is associated with increased mitochondrial biogenesis (Bhattacharya and Scimè, 2020). This is widely known for muscle differentiation, but whether dedifferentiation is marked by the reverse metabolic reprogramming has not been explored. IPA predicted activation of OXPHOS constituent pathways in the 5d differentiated A1 myotube state, as well as upregulation of mitochondrial biogenesis during differentiation and downregulation during cell cycle re-entry (Fig. 3D). These predictions were supported by expression patterns of TCA cycle genes and the free coenzyme A/Acetyl-CoA ratio, which is consistent with increased OXPHOS usage in myotubes (Fig. 3E and 3F, respectively). Whilst the electron transport chain gene set showed an increase at 48 hours of dedifferentiation (Fig. 3G), the available metabolic data is likely more reliable, and additionally supported by IPA analysis of mitochondrial biogenesis (Fig. 3D). We further profiled the dynamics of glycolysis rate, revealing downregulation of glycolytic enzymes lactate dehydrogenase (LDHA) and the muscle isozyme of pyruvate kinase (PKM) during differentiation, followed by upregulation during dedifferentiation (Fig. 3E&F). These changes were corroborated by reduced production of lactate during differentiation (Fig. 3G), used as a proxy for anaerobic glycolysis rate.

Overall, this analysis charts the temporal dynamics of pathway activity and metabolic changes during A1 myotube differentiation and dedifferentiation, validating prior findings, highlighting new candidate drivers of such processes, and providing insights into cell fate-associated metabolic reprogramming.

### Hypoxia inhibits newt myotube dedifferentiation

To uncover early regulators of dedifferentiation, we compared the 8h dedifferentiating group to the 5d differentiated myotubes (Supplementary Fig. 2A). Among the top 20 most significantly differentially expressed genes, notable findings included the expected downregulation of the tumour suppressor p21 (CDKN1B, LFC = −2.3), a key factor required for cell cycle exit and a transcriptional target of p53 (Bodzak et al., 2008), consistent with previous findings (Yun et al., 2014). The most significant gene identified (padj = 8.1e-120, LFC = 3.1) was Refilin-B (Rflnb), a regulator of filamin B’s actin-binding properties during skeletal development and a downstream target of TGFβ (Baudier et al., 2018). The highest upregulated gene was tissue factor (TF/F3, LFC = 6.1), a protein essential for thrombin formation during blood clotting. TF has been shown to be activated by TGFβ in combination with serum, entering a positive feedback loop (Felts et al., 1997; Grover and Mackman, 2018; Shi et al., 2024). We also observed stark upregulation of protease inhibitors SERPINE1 and TIMP3, both well-established targets of TGFβ during ECM remodelling (Kong et al., 2021; Leivonen et al., 2013). Interleukin 11 (Il11) was another noteworthy hit, with a significant but transient overexpression peak at 8h of dedifferentiation (LFC = 5.1), returning to baseline after 48 hours. Interleukin 11 has recently been linked to limited fibrotic scarring during tissue regeneration in zebrafish (Allanki et al., 2021). Regulator of Cell Cycle (Rgcc), another suppressor of fibrotic signalling activated by chemically-induced dedifferentiation in myofibroblasts (Fortier et al., 2021; Penke et al., 2021; Tian et al., 2023), was upregulated in a similar pattern (LFC = 4.3).

To profile broader temporal patterns, we analysed all significantly differentially expressed genes in the 128h vs 120h comparison by clustering their temporal profiles and running enrichment analysis on the clusters. However, the results were largely nonspecific, with processes such as cell cycle regulation, DNA replication, ECM remodelling, wound healing, and developmental processes being highlighted – all terms expected from cells changing their identity (Supplementary Fig. 2B-D). Notably, response to TGFβ also appear within the gene enrichment analysis cluster 8, showing strong transient upregulation during dedifferentiation.

Yet, the most compelling among individual changes was the upregulation of Hypoxia Inducible Factor 3 Subunit Alpha (HIF3A) (LFC = 3.77; Fig. 4A-C), as its effect on dedifferentiation had not yet been described. Profiling of other hypoxia-related genes revealed no significant changes in expression of HIF1A or its inhibitor HIF1AN. However, since HIF1A is primarily post-transcriptionally regulated, the expression of glycolytic enzymes shown in Figure 3H&I serves as a more reliable marker of upstream HIF1A activity (Semenza et al., 1994). In contrast, HIF2A/EPAS and HIF1B/ARNT were significantly upregulated during differentiation and downregulated during dedifferentiation (Fig. 4B, Supplementary Fig. 3E&G), along with corresponding expression changes in HIF2A targets SOX2, BHLHE40 and SPRY1. Interestingly, HIF-degrading prolyl hydroxylase domain protein PHD1/EGLN2 (Appelhoff et al., 2004) showed a trend resembling the transient HIF3A upregulation during dedifferentiation (Supplementary Fig. 3F). Further, IPA predicted upregulation of cellular response to hypoxia in myotubes and downregulation during dedifferentiation, consistent with the observed HIF2A and EGLN2 expression changes (Fig. 4D).

**Figure 4.**
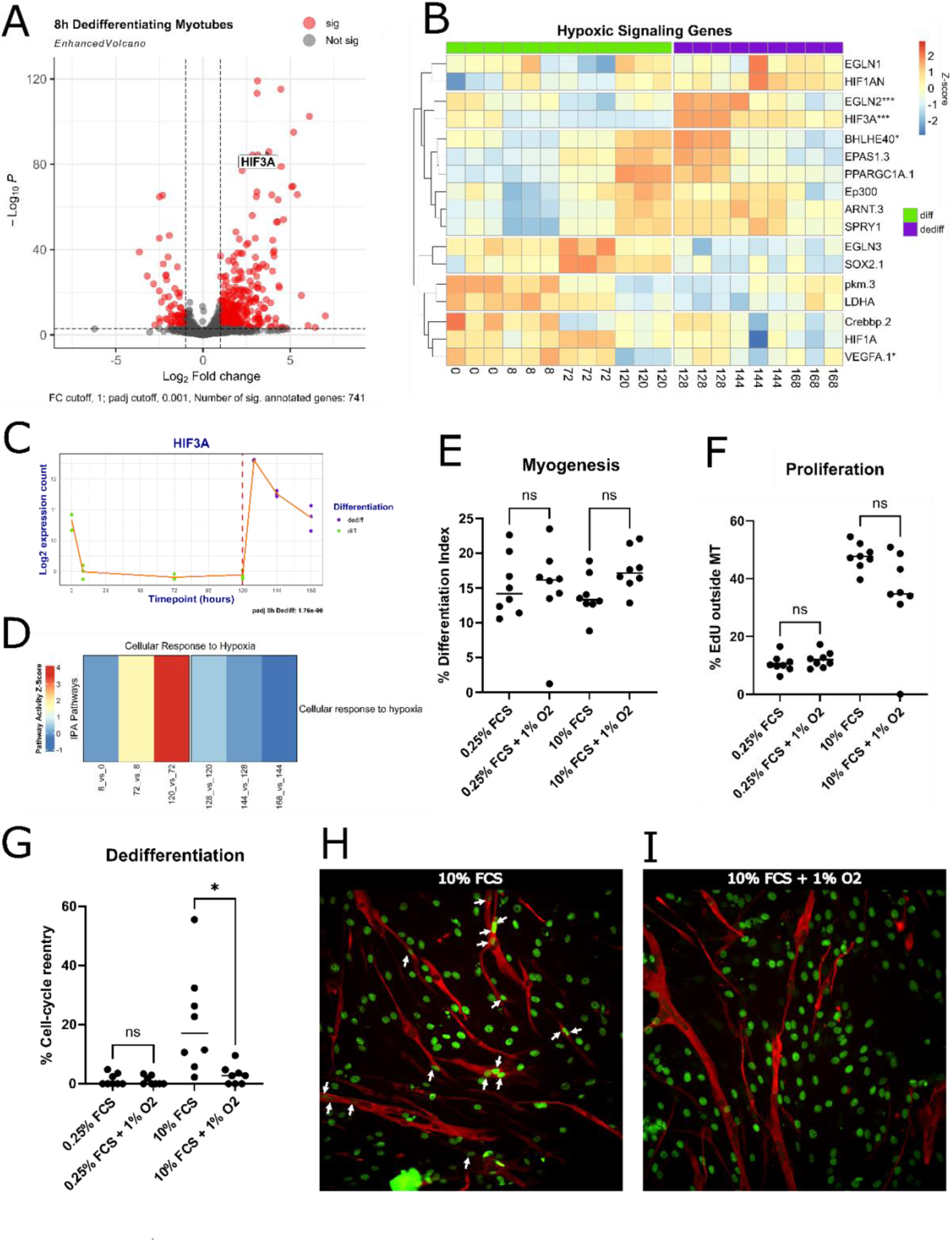
Hypoxia signalling inhibits cell-cycle re-entry of A1 myotubes. A) Volcano plot highlighting hypoxia-inducible factor 3 alpha (HIF3A) among the most prominent genes upregulated in early dedifferentiation. Significant annotated genes highlighted in red with log2fold change cutoff of 1 and padj cutoff < 0.001. B) Clustered heatmap showing member genes of hypoxic signalling. Time points are arranged in columns in chronological order, vertical cut and colour legend indicate induction of dedifferentiation. Stars on rownames indicate significance padj < 0.001, after a Wald test comparing 128h to 120h (early dedifferentiation; *p < 0.05, **p < 0.01 and ***p < 0.001). C) Gene expression changes of HIF3A throughout the differentiation and dedifferentiation time course. Y-axis: log2-transformed normalized counts. X-axis - time point in hours. The orange line connects means of log2 counts per time point. Red dotted line and colour legend indicate induction of dedifferentiation. Adjusted pvalue shown after a Wald test comparing 128h to 120h. D) Heatmap of cellular response to hypoxia IPA pathway. Each cell represents a consecutive comparison to the previous timepoint, arranged in columns. Z-scores were taken directly from IPA. E-I) Quantification results of dedifferentiation under normoxic and hypoxic conditions, with E) showing % of nuclei in myotubes, F) % EdU-positive proliferating cells; and G) % EdU positive nuclei in myotubes. Significance stars shown for Kruskal-Wallis test with Dunn’s post-hoc pairwise comparisons. H, I) Representative images from A1 cell-cycle re-entry assay under normoxic and hypoxic conditions respectively. Green - EdU, red - myosin heavy chain staining. White arrows point to EdU-positive nuclei in myotubes.

Based on current evidence, A1 myoblasts would be expected to exist in a hypoxic state to preserve their stemness in the face of shifting metabolic demands during differentiation (Elashry et al., 2022). This is consistent with our metabolic data (Fig. 3D-J), but contrasts with the observed transcriptomic changes. To address this inconsistency, we investigated the effect of hypoxia on both myogenesis and dedifferentiation, by exposing cells to low or high serum medium in 1% O2 (hypoxia) or 21% O2 (normoxia). The results showed that hypoxia had no effect on myogenesis rate regardless of serum presence (Fig. 4E), nor a significant impact on the proportion of dividing mononucleate cells in the 10% FCS group (Fig. 4F). Surprisingly, there was almost a complete block of cell cycle re-entry in hypoxic myotubes (∼18% reduction), with a re-entry rate comparable to that of serum-deprived groups (Fig. 4G, H, I). The experiment was repeated independently, whereby inhibition of dedifferentiation by hypoxia remained robust and replicable, with a sharp decrease in cell cycle re-entry (Supplementary Fig. 3H).

Altogether, our study demonstrates that serum-induced dedifferentiation is characterized by significant transcriptional changes, notably the transient upregulation of HIF3A and TGFβ targets. Moreover, we uncover that hypoxia inhibits myotube cell cycle re-entry, indicating that oxygen levels are critical modulators of myotube plasticity, a central process for the regeneration of complex structures.

## DISCUSSION

Here, we validated the myogenic A1 newt line as a physiologically relevant model, confirming its homogeneous myoblast identity. Through time-resolved transcriptomic profiling of newt myotube differentiation and dedifferentiation, we demonstrate that TGFβ signalling appears to be central to both cell cycle exit of A1 cells and the re-entry of A1-derived myotubes. We define the metabolic signatures that characterise each cell state, and uncover hypoxia strongly inhibits cell cycle re-entry in myotubes specifically, and identify HIF3A and HIF2 as putative mechanistic links between oxygen levels and induction of dedifferentiation.

Clarifying the cell identity and heterogeneity of the A1 line (Fig. 1) was important, as the line had never been purified yet extensively used for cell plasticity studies since its generation (Ferretti and Brockes, 1988; Lo et al., 1993). The finding that A1 cells constitute myoblasts suggests the existence of such population within newt limb tissues and thus raises questions on their origin, namely whether they derive from -as yet elusive-satellite cells, or whether they emerge from dedifferentiation of muscle fibres. These should be elucidated by future research.

The differentiation and dedifferentiation trajectory is evident in the PCA of our transcriptomic time course, showing a gradual adoption of skeletal muscle markers (Fig. 2C&F). Interestingly, dedifferentiating myotubes diverge from control cycling cells along the cell cycle-associated PC2 dimension, indicating that re-entry does not represent a simple return to the baseline proliferation state but rather reflects a new state with a distinct transcriptional profile.

While confirming previous observations of BMP and p53 signalling as indispensable for regulating myotube cell cycle re-entry (Wagner et al., 2017; Walters et al., 2023; Yun et al., 2013), TGFβ emerges as a key player in early dedifferentiation induction (Fig. 3A-D). This aligns with existing literature in mice showing that TGFβ inhibits myotube formation by repressing MyoD and reducing MHC expression via SMAD3 (Filvaroff et al., 1994; Liu et al., 2001), limiting mononucleate cell fusion (Girardi et al., 2021). Additionally, TGFB1 is upregulated during the healing and dedifferentiation phase, and downregulated during redifferentiation stage of axolotl limb regeneration, where blastema formation fails when TGFβ is inhibited (Lévesque et al., 2007). TGFβ also regulates developmental senescence via SMAD3 phosphorylation in axolotls and Xenopus (Davaapil et al., 2017), with senescence non-cell-autonomously promoting newt myotube dedifferentiation *in vitro* and limb regeneration *in vivo* (Walters et al., 2023; Q. Yu et al., 2022; Yun et al., 2015).

In the model used herewith, TGFβ is likely introduced directly with the serum, as its latent form is highly abundant in fetal serum (FBS/FCS) (Oida and Weiner, 2010). Latent TGFβ becomes active when cleaved from the latency-associated peptide (LAP) by proteases like thrombin, plasmin or matrix metalloproteinases (MMPs) to release active TGFβ (Annes et al., 2003; Blakytny et al., 2004; Chang et al., 2017; Jenkins, 2008). MMPs are known to be essential for newt limb regeneration (Vinarsky et al., 2005), and we observed an appropriate, transient spike of MMP expression during dedifferentiation (Supplementary Fig. 3C). Inhibition of protease activity using the broad-spectrum MMP inhibitor GM6001 or serum protease inhibitor AEBSF inhibits myotube cell cycle re-entry to a varying degree (Wagner et al., 2017; Walters et al., 2023). Notably, Il11 expression also coincides with MMP activity in early axolotl blastema formation (Gerber et al., 2018) and promotes cellular reprogramming while limiting profibrotic effects of TGFβ during zebrafish tissue regeneration via STAT3 (Allanki et al., 2021). Thus, the transient peak of Ill11 we detect during dedifferentiation induction (Supplementary Fig. 1E) likely serves towards a broader regulatory role in mitigating TGFβ overactivation.

Another relevant finding that is the noticeable upregulation of tissue factor (TF/F3) during early A1 dedifferentiation (Supplementary Fig. 1D). TF leads to the production of thrombin via a multi-phase process (Brummel et al., 2002; Cate et al., 1993; Wood et al., 2013), which plays a dual role in regeneration: initiating the coagulation cascade and proteolytically cleaving free-floating growth factors in serum. Thrombin activity is critical for salamander regeneration, in particular newt lens regeneration, which also involves regulation of cell plasticity and whereby TF localization coincides with the region of thrombin activation (Godwin et al., 2010; Imokawa and Brockes, 2003; Simon and Brockes, 2002). This is especially interesting in light of the study by Tanaka et al., 1999; showing that thrombin acts as a serin protease, cleaving dormant growth factors in serum into their active ligand forms. Indeed, Wagner et al., 2017 later characterized the re-entry promoting effects of thrombin-cleaved BMP ligands on newt myotubes, identifying BMP as one of the putative serum-supplied substrates of clotting factor proteases. However, the small size of TGFβ subunits and homodimers (Roberts et al., 1985, 1983) may have prevented detection in the previous work. It is thus possible that TGFβ could also constitute an early target of thrombin, relevant to muscle dedifferentiation.

Blocking BMP receptors abolishes myotube re-entry (Wagner et al., 2017; Walters et al., 2023). Given the temporal dynamics of pathway activation (Fig. 3A&B; Supplementary Fig. 3A&B), TGFβ may act either upstream of or synergistically with BMP, consistent with known cross-talk between the pathways (Guo and Wang, 2009; Wang et al., 2020). This aligns with the mechanism proposed by Wagner et al., 2017, where amputation-induced tissue damage and exposure to animal’s own plasma induce the regeneration response, triggering both the coagulation cascade as well as proteolytic cleavage of growth factors. Our findings suggest TGFβ as another key substrate of proteolytic cleavage by clotting factors that induce myotube dedifferentiation. Future studies should directly address whether transient TGFβ exposure can induce dedifferentiation in postmitotic myotubes.

The observed shift from anaerobic glycolysis in A1 myoblasts to OXPHOS in differentiated myotubes (Fig. 3D-J) aligns with evidence that metabolic reprogramming is required for myogenic differentiation (Lyons et al., 2004; Shintaku et al., 2016; Sin et al., 2015). Our findings suggest dedifferentiating myotubes reverse this process, adjusting energy metabolism from OXPHOS back to anaerobic glycolysis.

Metabolic reprogramming during muscle differentiation closely relates to oxygen availability and the hypoxic signalling system. High lactate production and hypoxia are hallmarks of stem cells and highly proliferative cells overall, widely described for cancer cells as the Warburg effect (Hanahan, 2022). In skeletal muscle, hypoxia promotes myoblast proliferation, preserves stemness and enhances Pax7+ stem cell proliferation in various mammalian models. Additionally, it blocks myogenic differentiation by downregulating MYOD1 and MYF5 and preventing cell cycle exit (Di Carlo et al., 2004; Elashry et al., 2022; Endo et al., 2024; Lees et al., 2008; Pircher et al., 2021; Urbani et al., 2012).

HIF1A favours a metabolic shift towards aerobic glycolysis in proliferative cells (Lindholm and Rundqvist, 2016; Taylor and Scholz, 2022) by upregulating virtually all glycolytic enzymes, including PKM and LDHA, the corresponding changes of which we also detect (Fig. 3H&I) (Iyer et al., 1998; Luo et al., 2011; Samanta and Semenza, 2018). That response is unique to HIF1A, with the other HIFs having less understood, context-dependent functions (Franke et al., 2012; Lee et al., 2024; Loboda et al., 2010).

Our metabolic profiling indicates that A1-derived myotubes are not hypoxic and rely on OXPHOS more so than the undifferentiated A1 cells (Fig. 3D-J). It is thus counterintuitive to see overexpression of HIF3A during dedifferentiation, reported to have a dominant negative inhibitory function over the other HIFs via competitive binding to the HIFβ subunit (Makino et al., 2001; Ravenna et al., 2016). However, our transcriptomic data revealed that the effects are likely mediated via interfering with HIF2A, which is gradually downregulated along with its target genes and HIF1B, contrasted by a peak of overexpression of HIF-degrading prolyl hydroxylase ELGN2/PHD1 (Fig. 3B; Supplementary Fig. 3E-G). Thus, HIF3A likely acts to rapidly block HIF2A signalling before its downregulation can take place.

Yang et al., 2017 showed that HIF2α is expressed at higher levels in post-differentiation myotubes, where it supports myofiber function by upregulating SPRY1—a pattern also observed in our system (Fig. 4B). The effects appear to be connected to the maintenance of oxidative phosphorylation and enhance mitochondrial function. One of the mechanisms by which TGFβ reduces differentiation efficiency in myoblasts is by downregulating the master-regulator of mitochondrial biogenesis PGC1α (PPARGC1A), linking it to metabolic reprogramming (Hoffmann et al., 2018). The expression of PGC1α in our system correlates with that of HIF2A (Fig. 4B) and negatively correlates with TGFβ activity. PGC1α itself was shown to act upstream of HIF2A, supporting the oxidative muscle fibre phenotype (Rasbach et al., 2010). This suggest TGFβ may act upstream of HIF2A. The existence of a causal link between TGFβ and HIF2A signalling in muscle dedifferentiation merits further investigation. Future research should directly explore the potential of reducing HIF2A signalling in promoting newt muscle dedifferentiation, by overexpression of HIF3A or specific inhibition of HIF2A in myotubes.

## LIMITATIONS OF THE STUDY

Although we have early dedifferentiation data, HIF-mediated cellular responses can occur within the first hour of exposure. Introducing even higher resolution time points would thus provide better insight into time-dependent dynamics of dedifferentiation induction. The lack of a genome assembly for *N. viridescens* complicates the precise functional analysis of certain genes, particularly HIF3A, which has multiple splice variants with distinct functions. Additionally, since HIFs are largely regulated via posttranscriptional degradation, we cannot determine the stabilization status of the HIF proteins themselves as mere expression level may not reflect activity.

## MATERIALS AND METHODS

### Cell culture

A1 cells derived from *N. viridescens* newt limb (P. Ferretti and Brockes, 1988) were cultured as previously described (Oliveira et al., 2022; Yu et al., 2022); in brief, cells were grown on gelatin-coated plastic in MEM (Gibco) supplemented with 2 nM L-glutamine (Gibco), 10 μg/ml insulin (Sigma), 100 U/ml penicillin/streptomycin (Gibco), 10 % heat-inactivated FCS (Gibco) and 25 % dH2O. Cells were kept in culture at 25 °C and 2 % CO2 and split 1:2 when approaching 70-80 % confluence.

In all experiments the formation of myotubes (MT) was induced by culturing A1 cells in medium with 0.25 % FCS (PAA Laboratories) serum concentration. After 5 days MT formation was observed, and the media was exchanged for 10 % PAA FCS serum to promote cell cycle re-entry and dedifferentiation.

### Cell cycle re-entry assay

A1 cells were seeded on gelatin-coated 96-well plates (150 % confluence in total volume of 200 µl), differentiation and dedifferentiation were induced as described above. One hour before reaching the timepoint of interest the cells were exposed to 1 µl of 1 mM EdU (Invitrogen), for a final EdU concentration of 5 µM. When the timepoint was reached, the cells were fixed in 4 % cold paraformaldehyde (in PBS) for 10-15 min. After fixation, the cells were covered with A-PBS (80 % PBS, 20 % dH2O v/v) and stored at +4 °C until staining.

### Click-iT EdU and Immunostaining

Click-iT EdU staining was performed according to manufacturer’s instructions (Invitrogen); in brief, the fixed cells were permeabilized with 100 µl 0.2 % Triton-X100 in PBS for 30 min at room temperature (RT) and stained with a freshly made Click-iT reaction cocktail (Antibody: Alexa Fluor azide 488 or 594). Prior to immunostaining, samples were blocked in 10 % goat serum in PBS for 30 min (RT), and then stained with antibodies listed in Table 1. Primary antibodies were incubated overnight at 4 °C and secondary antibodies for 1-4 h at RT. Antibody solution was made up in 5 % goat serum and 0.1 % Triton-X100 in PBS, and samples were washed twice in PBS between primary and secondary incubation. Nuclei were counterstained using 1:2000 Hoechst 33342 (Invitrogen).

**Table 1.**
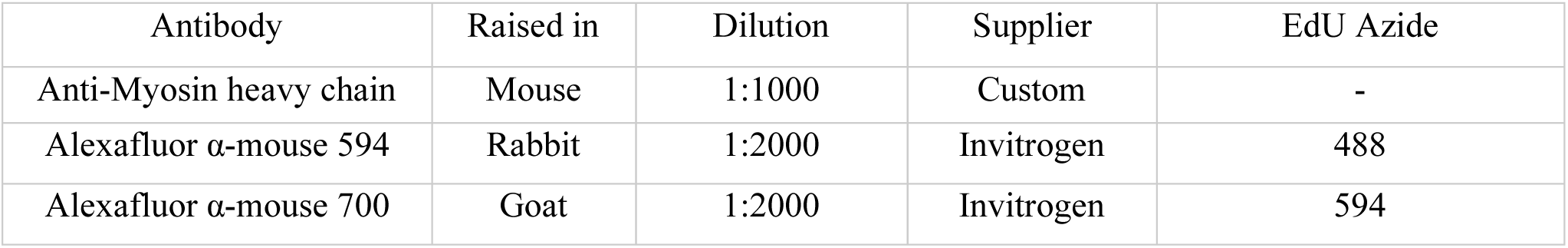
Antibodies used for Immunostaining.

| Antibody | Raised in | Dilution | Supplier | EdU Azide |
| --- | --- | --- | --- | --- |
| Anti-Myosin heavy chain | Mouse | 1:1000 | Custom | - |
| Alexafluor α-mouse 594 | Rabbit | 1:2000 | Invitrogen | 488 |
| Alexafluor α-mouse 700 | Goat | 1:2000 | Invitrogen | 594 |

### Imaging & Data analysis

Multichannel images of the stained cells were taken using a Nikon Eclipse TsER microscope. FIJI was used for blinded image analysis. For total nuclei number, watershed was applied to a binary image of Hoechst staining, and the nuclei were counter with FIJI’s Analyze Particles. EdU and nuclei in myotubes were counted manually. Data acquired from images was analyzed using Excel 2016 (Microsoft Corporation, Redmond, USA), and Graph Pad Prism 9.3.1 (San Diego, USA).

### Cell harvest and myotube purification

For bulk RNAseq, cells were cultured in either 30 mm^2^ (control) or 10 cm^2^ (other timepoints) dishes, scored with a scalpel for better MT formation, three biological replicates each. Differentiation and dedifferentiation were induced as described above. After reaching the timepoint of interest, cells or myotubes were washed once with A-PBS and then lifted by brief trypsinization (A-Trypsin made up in A-PBS), followed by quenching with fresh 10 % FCS media. Samples before 5d of differentiation were simply spun down, lysed in 75 µl RTL buffer (QIAGEN) and frozen at −80 °C until RNA extraction. Myotubes starting from 5 d of differentiation were filtered through a 100 µm sieve for aggregate exclusion. The flow-through was filtered again through a 35 µm mesh for single cell exclusion, with myotubes retained on the filter. The MT were then washed off with 10 % FCS medium, spun down, lysed in 75 µl RTL buffer and frozen at −80 °C until RNA extraction.

### HPLC analysis of CoASH and AcetylCoA

CoA compounds measurements from newt cell extracts were performed as previously (Tsuchiya et al., 2014). Briefly, CoASH and short-chain CoA esters were extracted in 5 % perchloric acid (PCA) and further processed for HPLC injection, including PCA neutralisation and adjustment to pH 6. HPLC was conducted at 40 ° C using a Kinetex C18 column (100 × 4.60 mm) with 2.6 mm particle size and 100 Å pore size (Phenomenex). The following solvents were used: solvent A (150 mM NaH2PO4 and 9% methanol), solvent B (150 mM NaH2PO4 and 30% methanol). CoA compounds were eluted isocratically with solvent A at a flow rate of 0.8 ml/min for the first 20 minutes after which a linear gradient to 100% solvent B was applied over 5 minutes at a flow rate of 0.5 ml/min. This was followed by a linear gradient back to 100% solvent A and the flow rate was returned to 0.8 ml/min over 5 minutes. Elution of CoA compounds was monitored by absorbance at 254 nm. Peaks corresponding to CoA compounds in newt cell extracts were identified by comparison of retention times with those of authentic standards determined on the same day. CoA compounds were quantified by comparison of peak areas with those of external standards. Peak area was quantified by Borwin chromatography software. CoA standards and all common chemicals were from SigmaAldrich.

### Enzymatic Lactate Assay

To measure lactate levels, A1 cells or myotubes were washed with Minimum Essential Medium (MEM) without serum and incubated with 1000 µL of fresh MEM (no serum) for 6 hours. At 0 hour and 6 hour time points, cells were terminated by adding drops of perchloric acid (PCA, 5 % final concentration). Dead cells were scraped off from the plate, the resulting sample collected in 1.5 mL microcentrifuge tubes and then centrifuged at 10,000 rpm for 5 min to remove debris. The supernatant was transferred to fresh tubes, frozen in liquid nitrogen, and stored at −80°C until further analysis. The PCA extracts were neutralized with triethanolamine (TEA) and potassium carbonate (K₂CO₃).

For the lactate assay, a reaction buffer was prepared by dissolving 7.5 g glycine, 5.2 g hydrazine sulfate, and 0.2 g EDTA in 50 mL of water. A reaction mixture was prepared by mixing 2.5 mL of the reaction buffer with 0.00822 g NAD⁺, adjusting the pH to 9.5 in a final volume of 5 mL. For the assay, 50 µL of neutralized PCA extract was incubated with 130 µL of the prepared reaction mix at 30 °C for 5 min. Baseline absorbance was measured at 340 nm over 10 minutes before adding 20 µL of lactate dehydrogenase (LDH; 60 µL LDH/390 µL H₂O) to initiate the reaction. Absorbance at 340 nm was recorded until the reaction reached an endpoint, with the difference in absorbance to the baseline corresponding to the lactate concentration.

### Transcriptome annotation

The N. viridescence transcriptome was acquired from Tsissios et al., 2023 and previously annotated with GO terms and protein domains via InterPro (Hunter et al., 2009). Data obtained from InterPro was upended with “_IP” in the final annotation table. We subjected that transcriptome to further rounds of annotation. The transcriptome was ran through TransDecoder (github.com/TransDecoder) using Galaxy (Afgan et al., 2018) with default settings to obtain predicted protein sequences. Protein sequences were then annotated on a per-transcript level with blastp against UniprotKB’s Swiss-prot database with a significant homology hit E-value cutoff 10^-10. Uniprot ID hits were mapped to gene names using mapUniProt from the UniProt.ws R package v2.42.0 (Carlson M, 2023), Uniprot ID’s that lacked a gene name were mapped to protein names. In order to enable a quality control step for single-cell data, mitochondrial genes were manually annotated by homology to *Ambystoma mexicanum* mitochondrial genes.

For the purpose of downstream analysis, the blastp annotation was then used to create a tx2gene mapping file merging isoforms under the same gene ID, using a custom R script available here. Isoforms were defined as transcripts with the same consecutive number (e.g. tx5, tx6, tx7) and the same homology hit. Both Uniprot and InterPro hits were used for isoform assignment, prioritizing whichever had the longer group size (e.g., if Description_IP is x x y x but Uniprot is z z z z, Uniprot was preferred).

The blastp-annotated table was then merged with tx2gene mapping file and the SYMBOL column was created that included either gene or protein name if available, and then gene ID for unannotated genes. The SYMBOL column was uniquified in case of duplicated IDs (same homology hit for non-consecutive transcripts), giving a greater integer suffix after the gene name to each additional ID (e.g., E2F1, E2F1.1, E2F1.2 etc.), resulting in the final annotation table.

Gene ontology terms obtained from InterPro were assembled into an org.Nviridescens.eg.db annotation package using AnnotationForge (Marc Carlson and Hervé Pagès, 2023). The final annotation package contains gene ID, transcript ID, Uniprot ID, descriptions from SwissProt and InterPro, as well as gene ontology terms from the three categories. The annotation is available for install here.

Note that a small number of genes crucial for muscle differentiation were absent in the Tsissios et al. reference, namely Myf5, Myog, Myod1 and Mrf4. However, during the analysis it was clear that this still constitutes a more complete reference than the alternative Sandberg transcriptome (Abdullayev et al., 2013) lacking Pax7, Acta2, Lum, Vim and many other genes relevant for this study. In the case of the four missing genes, the Z-scores from Fig. 2F were acquired by re-aligning the data to the Sandberg transcriptome.

An R script combining annotations into the final annotation file, that includes the transcript IDs for mitochondrial transcripts is available here.

### Bulk RNA sequencing

Total RNA was extracted from A1 cells or purified myotubes according to manufacturer’s protocol (RNeasy Micro, QIAGEN). The concentration of all samples was measured on NanoDrop spectrophotometer. Libraries were prepared under Nextera XT DNA Library Prep under a low-input protocol. Paired-end, 100 base pair Illumina bulkRNA sequencing was performed by the Dresden Concept Genome Center (DCGC).

The raw counts data acquired from the DCGC was pre-processed using useGalaxy.org (Afgan et al., 2018). The adapter sequences were first trimmed with TrimGalore (github.com/FelixKrueger/TrimGalore), quality control was performed with FastQC, and the reads were subsequently aligned to the *N. viridescens* transcriptome (see Transcriptome annotation) with salmon (Patro et al., 2017). The gene-level counts were imported with tximport (Soneson et al., 2015) using the previously generated tx2gene mapping file.

### Single-cell sequencing

Single-cell suspension was prepared by multiplex labelling cell lines from two different species of newt, *N. viridescens* and *Pleurodeles waltl*. The *P. waltl* data was not used in this work. The preparation of a single-cell suspension followed the “Single Cell Suspensions from Cultured Cell Lines for Single Cell RNA Sequencing” guidelines from 10X Genomics. Pwl3 *P. waltl* cells at passage 34 and 65 % confluence were detached with 0.25 % trypsin from a T75 flask. Cells were washed with A-PBS, detached with 0.25 % trypsin, and collected by centrifugation at 1000 rcf for 3 minutes at RT. The supernatant was aspirated, and the cell pellet was resuspended in 1 ml of A-MEM by pipetting. Then, an additional 2 ml of media was added, and the cell count was determined using a hemocytometer.

The cell suspension was transferred to a 2 ml Eppendorf Protein LoBind tube and centrifuged at 300 rcf for 3 minutes. The supernatant was carefully removed, and the pellet was gently resuspended by adding and pipetting 1 ml of A-PBS with 0.04 % BSA (Bovine Serum Albumin). The cells were centrifuged again at 300 rcf for 3 minutes, and the supernatant was removed. To resuspend the pellet, 500 μl of A-PBS with 0.04 % BSA was added and gently pipetted. The cell suspension was then passed through a 40 μm cell strainer to remove cell clumps. The cell concentration was determined using a hemocytometer, with the target concentration falling in the range of 0.2 to 2 × 10^6 cells per ml. Finally, the single-cell suspension was processed together with the A1 sample following the Cell Multiplexing Oligo Labelling protocol from 10X Genomics.

For each sample, RT A-PBS with 0.04 % BSA was added for a total volume of 1 ml, and gentle pipetting was performed to mix the solution. The cells were then centrifuged at 300 rcf for 5 min at RT. The supernatant was carefully removed without disturbing the pellet. The cells were resuspended in 100 μl of Cell Multiplexing Oligo (CMO 311 for Pwl3, CMO 312 for A1), mixed by pipetting 10-15 times and incubated 5 min at RT. After the incubation 1.9 ml of chilled A-PBS with 1% BSA was added to the sample. The cells were centrifuged at 300 rcf for 5 min at 4 °C. The supernatant was discarded without touching the bottom of the tube to prevent pellet disruption. The pellet was resuspended in 2 ml PBS with 1 % BSA and gently mixed by pipetting. The cells were then centrifuged at 300 rcf for 5 min at 4 °C, and the supernatant was removed without disturbing the pellet. The previous wash step was repeated once more and the cell pellet was resuspended in 200 ul of chilled PBS with 1 % BSA. The cells were gently mixed by pipetting 10-15 times. Cell concentration and viability of the pooled sample were determined using a hemocytometer and propidium iodide. The labelled samples were pooled in a 1:1 ratio, considering the calculated concentrations. The pooled sample, with a concentration within the range of 1000-2000 cells/μl, was utilized for loading the 10X chip.

Cell encapsulation, complementary DNA (cDNA) generation, preamplification as well as library preparation were performed using Chromium Next GEM Single Cell 3ʹ v3.1 Kits according to the manufacturer’s instructions. Briefly, the pooled sample of 22.000 cells from two species was loaded on single-cell G Chip, the cell lysis and reverse transcription were performed on the chip. Post-GEM cDNA clean-up was performed with DynaBeads MyOne Silane beads and the cDNA was amplified using the CellPlex protocol with 11 amplification cycles. Small (CellPlex barcodes) and large cDNA fractions were separated using 0.6x SPRIselect beads (Beckman Coulter). The quality of the amplified cDNA and fragment size distribution was analysed using Fragment Analyzer (Advanced Analytical Technologies). Sequencing libraries were generated using Dual Index Kit TT Set A (6 amplification cycles were used for small cDNA fraction, 8 for the large one), assessed by Fragment Analyzer and sequenced on Illumina Novaseq 6000 (paired-end, 100 bp).

The multiplexed data from two newt species was computationally separated by the Dresden Concept Genome Center (DCGC), the *P. waltl* data was discarded for this work. Reads were aligned to the transcriptome by DCGC using cellranger-7.1.0. The filtered feature matrix was imported into R and processed with Seurat v5 (Baysoy et al., 2023; Butler et al., 2018; Hao et al., 2021; Satija et al., 2015). Transcript-level counts were summarized to genes, and a Seurat object was created using the suggested cutoffs of min.cells = 3 and min.features = 200. The final object contained 2302 cells.

Cell cycle related differences were regressed out using newt orthologues from the Seurat-provided S / G2M gene list (Supplementary Fig. 1A). Plots were generated with either ggplot2 (Hadley Wickham, 2016) directly in R, or using visualizations from ShinyCell (Ouyang et al., 2021).

### Cell type characterization

To determine the cell type of A1 cells, we had to compare gene expression levels for genes within the same sample, accounting for both sequencing depth and transcript length. As transcript per million (TPM) values are the most robust measure for within-sample comparisons, bulk samples from A1 cells (this study) and C2C12 mouse myoblasts (GSE247438) were separately aligned with kallisto (Bray et al., 2016) to obtain the TPM values. Genes were sorted by the median TPM value across biological replicates to account for outliers. Median TPM values were log2(median_tpm + 1) transformed to enable comparison between genes with wide range of expression (Fig. 1D, E). Cell type specificity of gene markers was inferred by referencing published research, as well as using data for corresponding orthologues from the Human Protein Atlas (Karlsson et al., 2021). An R notebook for marker comparison is available here.

### Differential gene expression analysis

The gene-level raw “counts” from salmon were further used for differential gene expression analysis (DGE) with DESeq2 (Love et al., 2014) in RStudio (Posit team, 2023). As we were interested in early dedifferentiation, the main comparison of interest was 128 hours (8h dedifferentiation) vs 120 hours (5 days terminally differentiated myotubes). The DESeq dds object was created with design = ∼timepoint, setting 120h as the baseline. Raw counts were filtered to at least 10 counts in at least three samples (smallest group size). The results table was extracted using res <-lfcShrink(dds, coef = “timepoint_128_vs_120”, type = “apeglm”) to shrink inflated log fold changes using apeglm (Zhu et al., 2019).

Normalized count data for individual gene plots was extracted from the dds object using DESeq’s plotCounts and then modified with ggplot2. EnhancedVolcano (Blighe K, Rana S, Lewis M, 2023) was used for volcano plots. Variance-stabilized transformed data vst(dds) was used downstream for PCA, clustering and heatmaps using pheatmap (Raivo Kolde, 2018).

For functional GO enrichment analysis, the significant genes obtained from the 128h_vs_120h comparison were filtered by Benjamini-Hochberg adjusted value of < 0.001. The data for those genes were then extracted from the vsd matrix for the entire differentiation / dedifferentiation time course. The values were centred and scaled to acquire normalized Z-scores, which were then clustered using k-means clustering (Supplementary Fig. 2D, E). An elbow plot was used to determine the initial number of clusters. Enrichment analysis was performed with ClusterProfiler (Wu et al., 2021) using the org.Nviridescens.eg.db annotation package generated above, taking only GO-annotated genes from the transcriptome as background. Clusters were compared using the enrichGO function with Biological Process (BP GO) ontology, with a pvalue cutoff of <0.05. Redundant terms were simplified with clusterProfiler::simplify cutoff = 0.7. Cnetplot exploring cluster overlap was generated using cnetplot().

Gene set plots were generated with a custom function, pulling vst-transformed values of all genes annotated with a given GO term. The values were centred and scaled, and means of the resulting Z-scores were plotted per timepoint. Lines connecting the means are shown on the final gene set plots on Fig. 2C and 3E&G.

The entire DEG script including custom plotting functions and enrichment analysis is available as an R notebook here.

### Ingenuity pathway analysis

LFC and padj data for Ingenuity Pathway Analysis (QIAGEN IPA, (Krämer et al., 2014)) was acquired by running DESeq2 with apeglm LFC shrinkage as described in the DGE section. Every timepoint was compared to the previous timepoint as the baseline, to profile high-resolution granular changes at each stage compared to the previous stage. Comparing each timepoint to 0h proliferating cells as the baseline was also trialed, however data was overwhelmed by nonspecific differentiation and cell cycle pathways and did not provide any useful information.

In IPA, both Uiprot ID and protein symbol columns acquired during annotation was used to map as many genes to the IPA database as possible. A core analysis was performed for each subsequent comparison, with absolute LFC cutoff of 0.5 and padj value cutoff of < 0.001. This combination again ensured that pathways with a smaller gene set size and milder changes would also be included in the analysis. A comparison analysis was created, and Z-scores for pathway or upstream regulator activation across all comparisons extracted into R. Heatmaps of pathway / upstream regulator activity were created using ComplexHeatmap (Gu, 2022), replacing N/A with zero (no change detected). Graphical summaries for 8h_vs_0h and 128h_vs_120h was extracted directly from IPA and shown on Supplementary Fig. 1B&C.

R notebooks for IPA data preparation are available here.

### Hypoxia induction

To ensure sufficient myotube formation in case of low effect size, myotubes were first formed for 6 days in normoxic conditions, and were then moved to the Whitley H35 Hypoxystation the Medizinisch Theoretisches Zentrum (MTZ) Dresden under 1% oxygen, adjusted to salamander culture conditions of 25°C and 2 % CO2. Normoxic control plates and hypoxic experimental plates were all seeded at the same time, from the same cell passage, with equal densities as described above. EdU pulse and fixing with PFA was done directly in the hypoxistation, maintaining 1 % oxygen levels.

## AUTHOR CONTRIBUTIONS

GV, HW, KT, YT and MHY designed experiments. GV, HW, KT, YT and MHY performed experiments. GV conducted all bioinformatic analysis. HEW conceived and supervised bulk transcriptomics experiments. KET performed single-cell sequencing. YT conducted metabolic assays. All authors interpreted data. GV wrote the first draft of the manuscript. MHY edited the manuscript with input from all authors. HEW and MHY acquired funding. MHY supervised the study.

## ACKNOWLEDGEMENTS

We thank Ben Wielockx for providing access to the Hypoxistation, advising on hypoxia experimental planning and data interpretation and Dresden Concept Genome Centre (CRTD, TU Dresden) for performing RNAseq. GV was supported by “Go West” scholarship from the Klaus-Tschira Foundation (KTS). KT was supported by a DAAD Scholarship. HEW was supported by an Alexander von Humboldt Stiftung post-doctoral fellowship. MHY was supported by Deutsche Forschungsgemeinschaft grants (DFG 22137416, 450807335 and 497658823) and TUD-CRTD core funds.

## ADDITIONAL INFORMATION

Supplementary Figures S1 - S3

