## Supplemental material for "Hypoxia inhibits newt skeletal muscle dedifferentiation"

ADDITIONAL INFORMATION

Supplementary Figures S1 - S3

FIGURE LEGENDS

**Supplementary Figure 1.**

Supplementary Figure 1. A) PCA plot, showing A1 single cell data without (left) and with (right) cell cycle regression. Colours indicate phases of the cell cycle. B, C) Graphical summary of top upstream regulators from IPA, for the 8h_vs_0h (B) and 128h_vs_120h (C) comparisons, highlighting TGFB1 as a central regulator for both processes. Cutoffs for IPA were 0.5 for LFC, p < 0.001, padj < 0.05. Orange - predicted activation, blue - predicted inhibition. D) Heatmap of top 20 IPA-predicted Upstream Regulators (UR) for the entire differentiation / dedifferentiation time course, in order of significance. Each cell represents a consecutive comparison to the previous timepoint, arranged in columns. Z-scores were taken directly from IPA. E) The same volcano plot as in Fig. 4A, all genes highlighted. Significant annotated genes highlighted in red with log2fold change cutoff of 1 and padj cutoff < 0.001.

**Supplementary Figure 2.**

Supplementary Figure 2. Transcriptomic exploration of early dedifferentiation induction of A1 myotubes. A) Clustered heatmap of top 20 most significant unique genes of 128h_vs_120h dedifferentiation. Time points are arranged in columns in chronological order, vertical cut and colour legend indicate induction of dedifferentiation. Stars on rownames indicate significance padj < 0.001, after a Wald test comparing 128h to 120h (early dedifferentiation; *p < 0.05, **p < 0.01 and ***p < 0.001). B) Cnetplot showing member genes by ontology, cluster and their overlap. Colour indicates cluster membership, size of dots represents amount of genes. C) Average progression of each cluster after k-means clustering of significant genes from the 128h_vs_120h comparison with padj < 0.001 cutoff. Y-axis - average Z-scores of all cluster member genes. X-axis - time point in hours. D) Biological Process Gene Ontology (GO BP) enrichment analysis across all clusters, showing shared terms where applicable. GO terms are arranged in rows, clusters in columns. Number of genes per cluster indicated next to cluster name. Dot size represents ratio of genes from total set size. Colour represents significance.

**Supplementary Figure 3.**

Supplementary Figure 3. Individual changes during differentiation and dedifferentiation of A1 myotubes. A, B) Clustered heatmap of ligands and SMAD signalling genes for BMP (A) and TGFβ (B). Time points are arranged in columns in chronological order, vertical cut and colour legend indicate induction of dedifferentiation. Stars on rownames indicate significance padj < 0.001, after a Wald test comparing 128h to 120h (early dedifferentiation; *p < 0.05, **p < 0.01 and ***p < 0.001). C) A plot showing overlapping Z-scores of SMAD1 (BMP) and SMAD3 (TGFβ). Lines are connecting average expression Z-scores (Y-axis) per time point (X-axis). D) Gene set plot, connecting the average expression Z-scores of all (104) genes annotated with Metalloendopeptidase Activity (GO:0008237) gene ontology term. E, F, G) Gene expression changes of HIF2A, PHD1 and HIF1B respectively throughout the differentiation and dedifferentiation time course. Y-axis: log2-transformed normalized counts. X-axis - time point in hours. The orange line connects means of log2 counts per time point. Red dotted line and colour legend indicate induction of dedifferentiation. Adjusted pvalues shown after a Wald test comparing 120h to 0h. H) Quantification results of dedifferentiation under normoxic and hypoxic conditions of an independent technical replicate, showing % of EdU positive nuclei in myotubes. Significance stars shown for Mann-Whitney test.

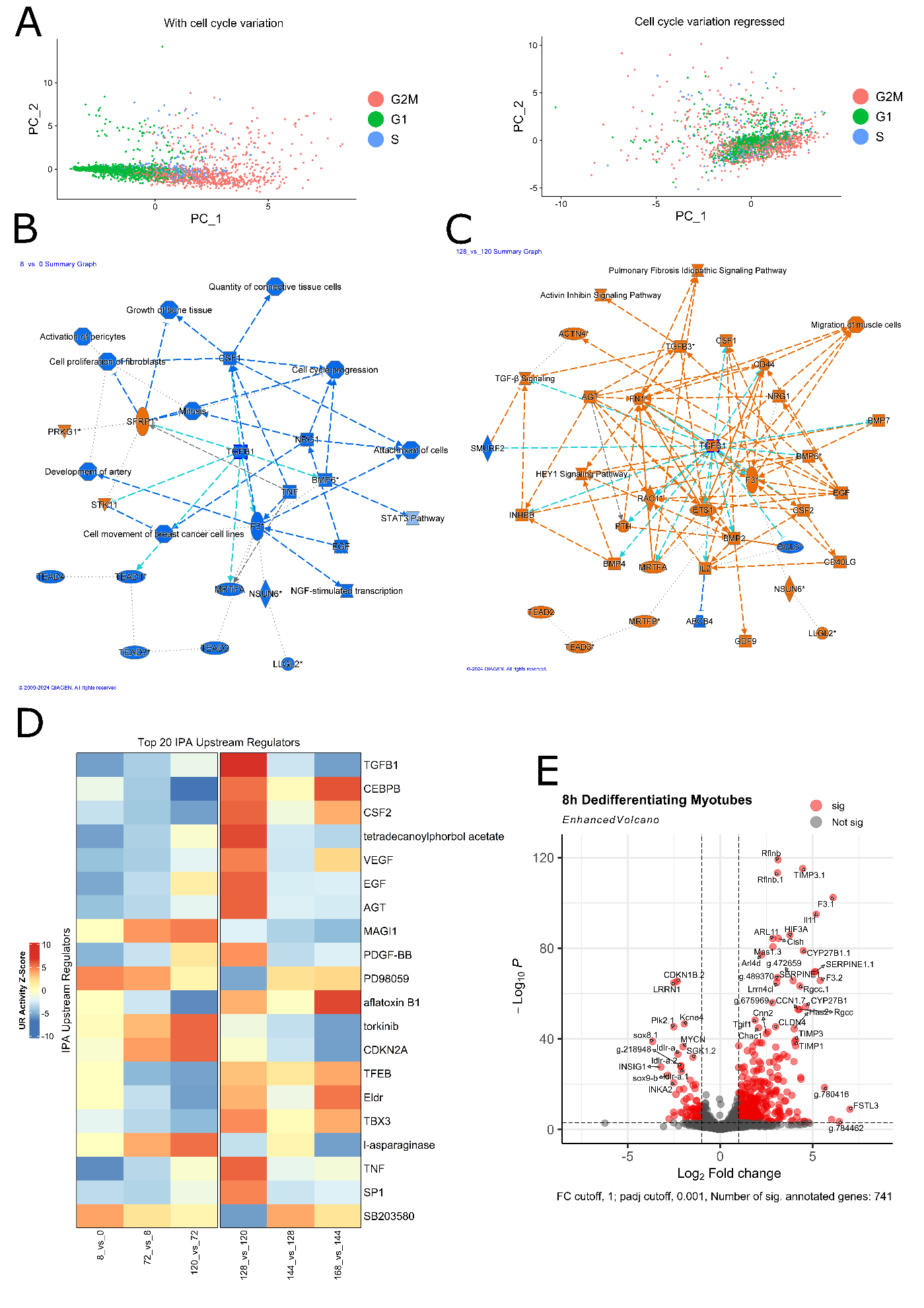

Supplementary Figure 1

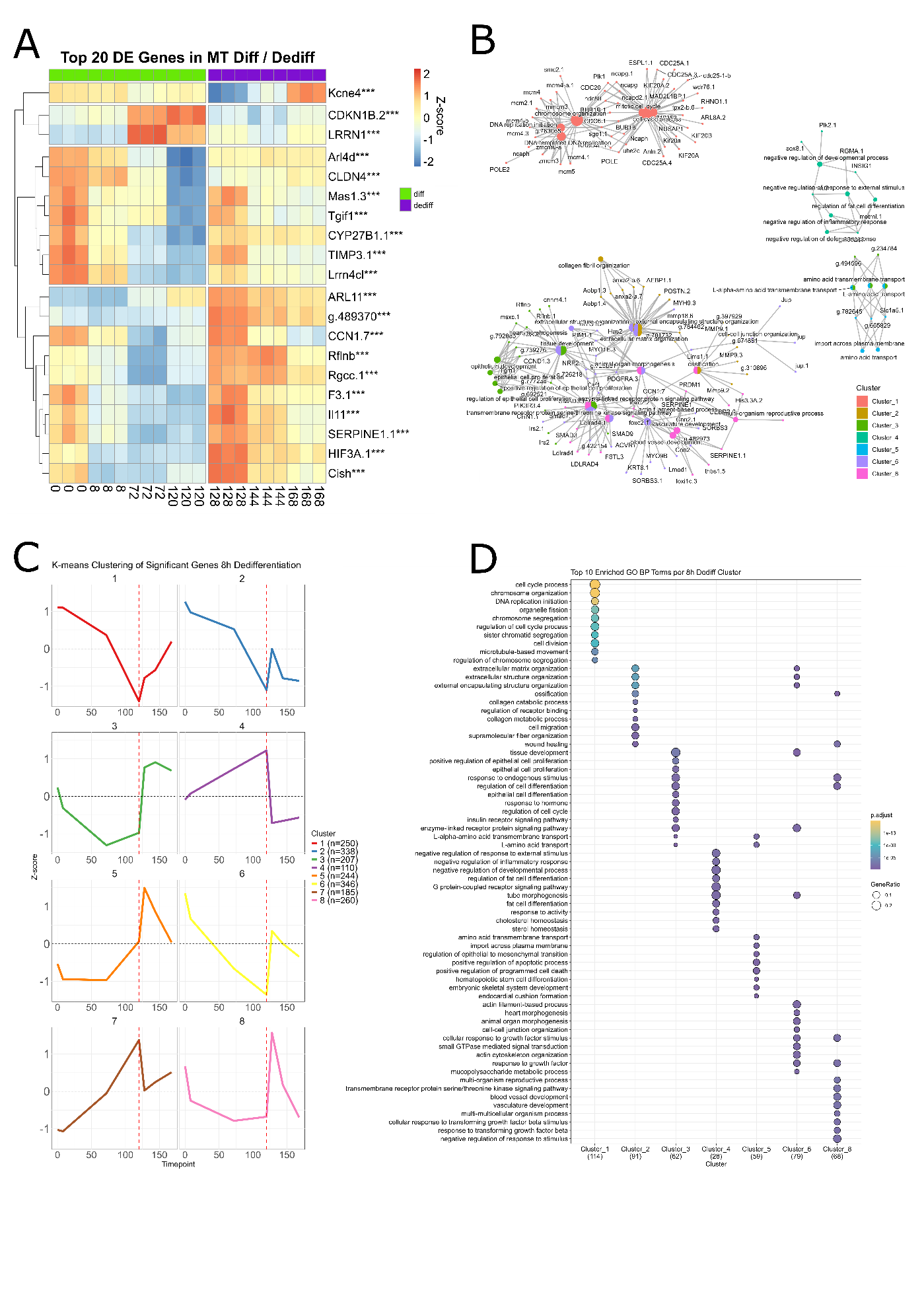

Supplementary Figure 2

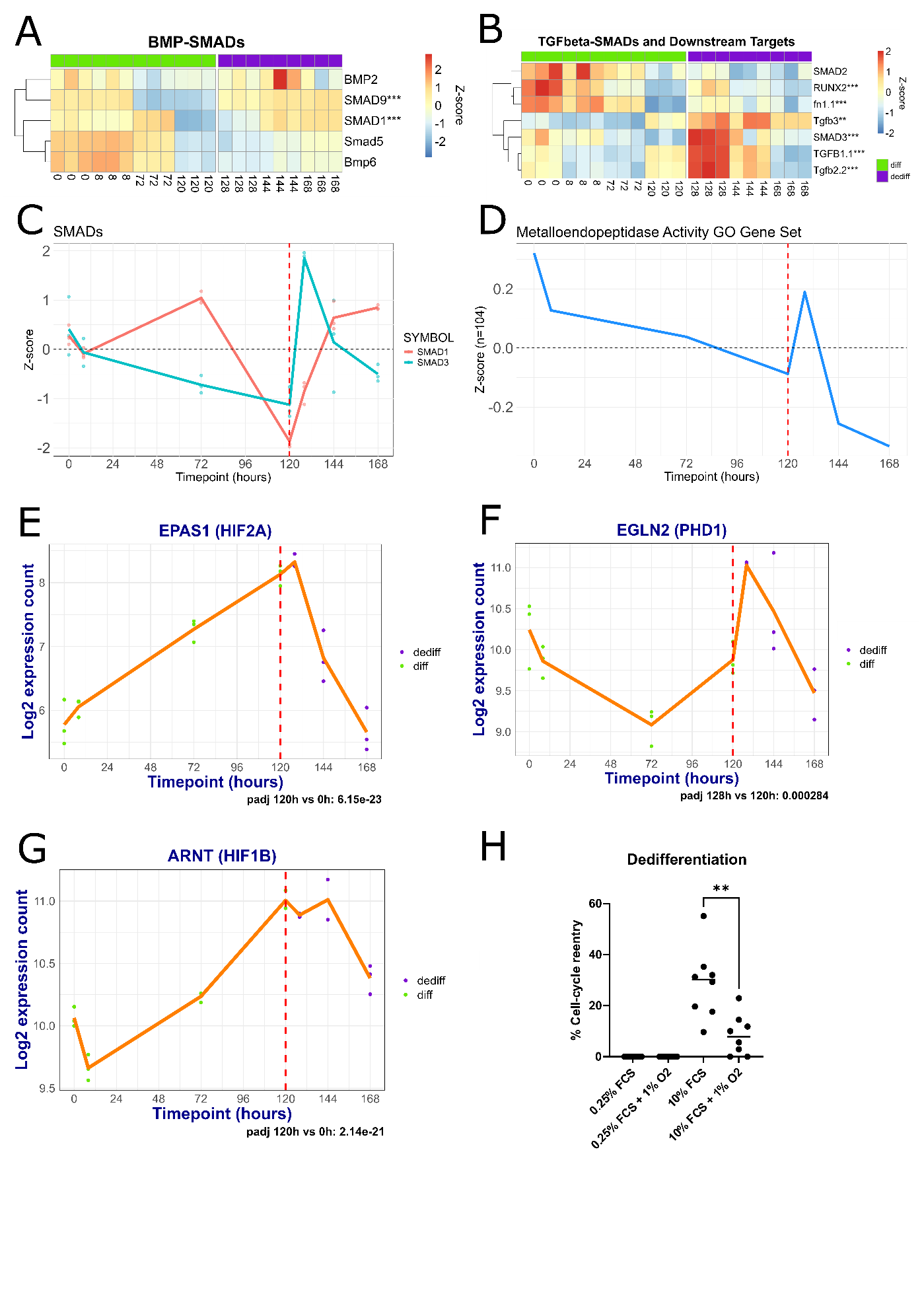

Supplementary Figure 3
